# Beyond Histology: A Validated CUBIC-Based Workflow for Volumetric Analysis of Follicles and Cortical Vasculature in Human Ovarian Tissue

**DOI:** 10.64898/2026.04.16.718954

**Authors:** Despina I. Pavlidis, Chloe E. Fischer, Maria A. Jennings, Jordan H. Machlin, Victoria Jan, Brendon M. Baker, Ariella Shikanov

**Affiliations:** Department of Biomedical Engineering, University of Michigan, Ann Arbor, MI, 48109, United States; Cellular and Molecular Biology Program, University of Michigan, Ann Arbor, MI, 48109, United States; Department of Obstetrics and Gynecology, University of Michigan, Ann Arbor, MI, 48109, United States; Department of Macromolecular Science and Engineering, University of Michigan, Ann Arbor, MI, 48109, United States

**Keywords:** Confocal microscopy, Human reproductive tissue, Light sheet microscopy, Ovarian tissue transplantation, Tissue clearing, Vascular architecture

## Abstract

**Research question:** Can tissue clearing, combined with volumetric imaging, enable reliable, quantitative three-dimensional analysis of follicles and vasculature in intact human ovarian tissue?

**Design:** A CUBIC-based clearing protocol was adapted for human ovarian medulla and cryopreserved cortex. Tissue from reproductive-aged donors was cleared, fluorescently labeled, and imaged using confocal and light sheet microscopy. Tissue expansion, imaging depth, and vascular morphometrics were quantified and follicle density was compared to conventional histology.

**Results:** Clearing produced optically transparent tissue with a linear expansion factor of 1.2 across cortex and medulla. Imaging depth increased 6.5–11-fold in cortex and 6–8-fold in medulla. Follicle density measurements in immunolabeled cleared cortex were comparable to histology, supporting the validity of volumetric follicle quantification. Light sheet microscopy of lectin-labeled cortex revealed no significant donor-to-donor differences in vascular morphometrics, including mean vessel diameters of 12–14 µm, branch point densities of 632–965 points/mm^3^, vessel length densities of 117–175 mm/mm^3^, and volume fractions of 1.9–2.3%. Volumetric imaging further illustrated heterogeneous spatial relationships between follicles and surrounding vessels.

**Conclusion:** Tissue clearing and volumetric imaging complement routine histology and enable quantitative three-dimensional investigation of follicle-vascular interactions in intact human ovarian tissue, providing a framework for advancing fertility preservation and ovarian tissue transplantation research.

## INTRODUCTION

Reproductive lifespan and long-term endocrine health depend on the finite ovarian follicle reserve and the complex stromal environment that supports this reserve. Together, the ovarian cortex, which is enriched in primordial and early growing follicles, and the ovarian medulla, which is more vascularized and houses larger developing follicles, constitute a spatially heterogeneous organ whose function is inseparable from its three-dimensional architecture. Follicles across both regions are maintained by vasculature, nerves, immune cells, and diverse stromal cell populations embedded in an extracellular matrix^1^. Although these components are well described in two-dimensional serial histological sections, their three-dimensional organization within intact human ovarian tissue remains incompletely characterized.

This limitation reflects the field’s long-standing reliance on section-based methodologies. While histology remains indispensable for evaluating follicle morphology and tissue composition, it reduces complex three-dimensional structures to two-dimensional snapshots, limiting reconstruction of spatial relationships across intact tissue volumes. Emerging spatial transcriptomic approaches have begun to reveal the cellular composition of the ovary within its spatial context^2–4^. These approaches, however, remain similarly constrained to thin tissue sections, underscoring the need for methods that preserve intact three-dimensional ovarian tissue architecture.

Tissue clearing offers a strategy to meet this need by rendering tissue optically transparent via reduction of light scattering, thereby enabling three-dimensional fluorescence imaging of large tissue volumes and even whole organs. Clearing has been widely adopted to investigate ovarian biology in mice, including follicle dynamics and vascular organization^5–12^. Translation to human ovarian tissue, however, requires validation as differences in stromal density, follicle distribution, and tissue scale^13^ can alter clearing efficiency, antibody penetration, and achievable imaging depth. A few recent reports have applied tissue clearing to human ovarian tissue, including characterization of the stromal proteome^14^, three-dimensional follicle visualization^15^, and comparative analysis of ovarian aging across species^16^. In this report, tissue clearing is extended to a standardized, validated workflow integrating clearing, fluorescent labeling, and quantitative volumetric analysis to support clinically relevant three-dimensional assessment of human ovarian tissue. This workflow is particularly relevant for fertility preservation strategies, such as ovarian tissue cryopreservation and transplantation, where preservation of the follicle reserve and timely vascular integration influence graft viability and endocrine function restoration^17^. Since follicle survival and vascular remodeling occur within a structurally complex stromal environment, quantitative three-dimensional mapping of follicles and surrounding vasculature and stromal compartments in intact ovarian tissue may strengthen graft outcome analyses and guide bioengineering strategies aimed at improving transplantation outcomes.

Here, we present a reproducible tissue clearing and volumetric imaging workflow for cryopreserved human ovarian tissue. Immunolabeling is implemented for follicle detection in cleared cortex, enabling volumetric follicle density quantification and benchmarking against conventional histology. Light sheet microscopy is subsequently applied to quantify vascular morphometrics in intact, lectin-labeled cortical tissue and to illustrate the spatial proximity between follicles and adjacent vasculature. Finally, confocal and light sheet imaging modalities are compared to provide practical considerations for implementation of our workflow. Together, this work establishes a reproducible, quantitatively validated platform that enables analyses not readily achievable with serial histological sectioning, directly addressing a methodological gap relevant to fertility preservation and ovarian tissue transplantation research.

## MATERIALS AND METHODS

### 1.1 Ethical Approval for the Procurement of Human Ovaries

As previously described^13,18^, whole human ovaries were obtained from deceased donors for non-clinical research through the International Institute for the Advancement of Medicine (IIAM) and associated Organ Procurement Organizations (OPOs). IIAM and the associated OPOs comply with state Uniform Anatomical Gift Acts (UAGA) and are certified and regulated by the Centers for Medicare and Medicaid Services (CMS). These OPOs are members of the Organ Procurement and Transplantation Network (OPTN) and the United Network for Organ Sharing (UNOS) and operate under standards established by the Association of Organ Procurement Organizations (AOPO) and UNOS. Informed, written consent was obtained from the deceased donors’ families prior to tissue procurement for the tissue used in this study. A biomaterial transfer agreement is in place between IIAM and the University of Michigan that restricts the use of the tissue to pre-clinical research not involving the fertilization of gametes. The use of deceased donor ovarian tissue in this research is categorized as “not regulated” per 45 CFR 46.102 and the “Common Rule” as it does not involve human subjects and therefore complies with the University of Michigan Institutional Review Board (IRB) requirements.

### 1.2 Collection of Human Ovarian Tissue

Human ovarian tissue was procured from four deceased donors (**Table 1**) who were of reproductive age and did not have a medical history indicative of conditions that would affect ovarian function. Upon arrival in the laboratory, donor ovaries were transferred to pre-cooled holding media composed of Quinn’s Advantage Medium with HEPES (QAMH, ART-1024, CooperSurgical, USA) supplemented with 10% Quinn’s Advantage Serum Protein Substitute (SPS, ART-3010, CooperSurgical, USA). The outer cortex layer was collected from the ovaries using a custom cutting guide (Reprolife, Japan) which produced tissue squares measuring approximately 1 mm in thickness and 10 mm in both length and width. Cortical squares were then cryopreserved using a previously published slow freezing protocol^13,18,19^. The remaining ovarian tissue, including the ovarian medulla, was fixed in 4% paraformaldehyde (PFA, J61899AK, Thermo Scientific Chemicals, USA) for 24 hours at 4°C. PFA-fixed tissue was then washed in Dulbecco’s phosphate-buffered saline (DPBS, 14190250, Gibco, USA) for 1 hour and stored in 70% ethanol (EtOH) at 4°C.

**Table 1.** Donor Metrics.

| **Donor** | **Age (years)** | **BMI (kg/m<sup>2</sup>)** | **CIT (hours)** |
| --- | --- | --- | --- |
| **A** | 31 | 37.7 | 27.7 |
| **B** | 28 | 22.7 | 20.4 |
| **C** | 31 | 24.2 | 29.9 |
| **D** | 34 | 24.4 | 10.5 |

BMI: body mass index; CIT: cold ischemic time

### 1.3 Preparation of Samples for Tissue Clearing

To prepare ovarian cortex for tissue clearing, cortex squares were thawed following a previously described protocol^18,19^. Briefly, cryovials containing cortex were removed from liquid nitrogen and left at room temperature for 30 seconds before being immersed in a 37°C water bath. Once the cryoprotectant medium had thawed (after approximately 1.5 minutes), the tissue was removed from the vial and placed into the first thawing solution (QAMH with 10% SPS, 0.5 M dimethyl sulfoxide (DMSO, D2650, Sigma-Aldrich, USA), 0.5 M Ethylene Glycol (102466, Sigma-Aldrich, USA), 0.1 M Sucrose (S1888, Sigma-Aldrich, USA)) for ten minutes. The tissue was then sequentially incubated in the second thawing solution (QAMH with 10% SPS, 0.25 M DMSO, 0.25 M Ethylene Glycol, 0.1 M Sucrose), the third thawing solution (QAMH with 10% SPS, 0.1 M Sucrose), and the fourth thawing solution (QAMH with 10% SPS) for ten minutes each. The fourth thawing solution served as a maintenance solution. All four solutions were equilibrated to room temperature for the thawing procedure. Thawed cortex squares were then manually cut into strips measuring approximately 5 mm in length and 1 mm in width. Cortex strips were subsequently fixed in 4% PFA for 24 hours at 4°C and then washed three times in DPBS for 1 hour each prior to being moved into 70% EtOH at 4°C for long-term storage. Medullary tissue was dissected from the fixed ovary remains and then manually cut into smaller pieces prior to undergoing the tissue clearing process.

### 1.4 Clearing of Tissues with Modified CUBIC Clearing Protocol

The CUBIC tissue clearing protocol described by Matsumoto et al.^20^ was adapted to clear human ovarian cortex and medulla samples. The protocol can broadly be separated into three phases: (1) delipidation and decolorization, (2) staining, and (3) refractive index (RI) matching (**Table 2**). The delipidation and decolorization phase began by immersing the prepared fixed tissue samples in an equivalent volume mixture of dH_2_O and CUBIC-L (10% (wt/wt) N-butyldiethanolamine (B0725, TCI America, USA), 10% (wt/wt) Triton X-100 (X100, Sigma-Aldrich, USA), 80% (wt/wt) dH_2_O) with shaking at 37°C overnight. The next day, this mixture was replaced with fresh CUBIC-L. CUBIC-L was refreshed every 48 hours for a total CUBIC-L incubation duration of 10 days. At the end of 10 days, to stop the delipidation and decolorization process, samples were washed with 1X phosphate buffered saline (PBS) with shaking at room temperature for at least 2 hours. This washing step was repeated more than three times.

**Table 2.**
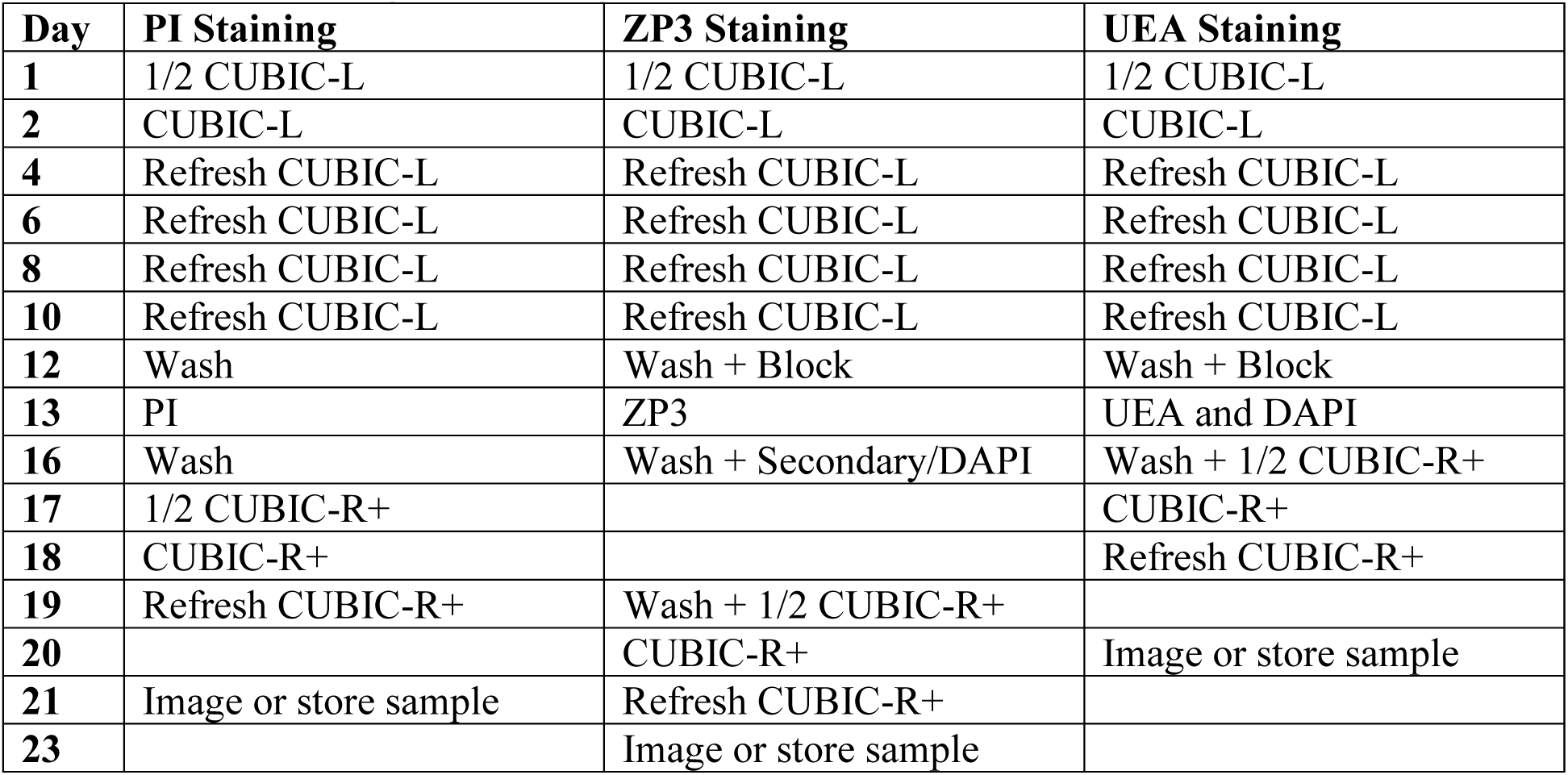
CUBIC Clearing and Staining Timelines.

Samples used for evaluation of imaging depth were stained with propidium iodide (PI, P3566, Molecular Probes, USA). To achieve this, the samples were immersed in staining buffer (1.5 M NaCl in 1X PBS) with shaking at room temperature twice for at least 2 hours each time. Then, the samples were moved to a staining buffer solution containing 30 μg/mL PI and were incubated for 3 days at 37°C with shaking. After the dye incubation step, the samples were washed with 1X PBS with shaking at room temperature overnight. To initiate the RI-matching phase of the CUBIC clearing protocol, the samples were then immersed in an equivalent volume mixture of dH_2_O and CUBIC-R+(M) (45% (wt/wt) antipyrine (D1876, TCI America, USA), 30% (wt/wt) N-methyl-nicotinamide (M0374, TCI America, USA), 0.5% (vol/vol) N-butyldiethanolamine, 24.5% (wt/wt) dH_2_O) with shaking at room temperature overnight. Next, this mixture was replaced with fresh CUBIC-R+(M) and the samples were incubated with shaking at room temperature overnight. The next day, CUBIC-R+(M) was refreshed and the samples were incubated for an additional day. After this final CUBIC-R+(M) incubation step, the samples were ready for imaging. The non-cleared samples did not undergo CUBIC-L and CUBIC-R incubation but were subjected to PI staining.

Samples used for evaluation of follicle density were stained using immunofluorescence labeling. After the CUBIC-L phase, the samples were immersed in blocking solution (5% normal goat serum (ab138478, Abcam, USA) in 1X PBS) with shaking at 37°C overnight. The samples were then incubated in a solution consisting of primary antibody (ZP3, 1:250, sc-398359, Santa Cruz Biotechnology, USA) diluted in blocking solution with shaking at 37°C for 3 days. Following this step, the samples were washed in blocking solution three times with shaking at room temperature for 1 hour each. Next, the samples were incubated in a solution containing secondary antibody (Goat anti-Mouse, Alexa Fluor Plus 647, 1:1000, A32728, Invitrogen, USA) and nuclear counterstain (DAPI, 1:1000, D1306, Invitrogen, USA) diluted in blocking solution with shaking at 37°C for 3 days. Finally, the samples were washed in 1X PBS three times with shaking at room temperature for 1 hour each before proceeding to the RI-matching phase described above.

Samples used for evaluation of vasculature were prepared using a modified version of the immunofluorescence labeling protocol described above. After the CUBIC-L phase, the samples were immersed in blocking solution (5% normal goat serum in 1X PBS) with shaking at 37°C overnight. The samples were then incubated in a solution consisting of Ulex Europaeus Agglutinin I (UEA I, DyLight 649, 1:250, DL-1068-1, Vector Laboratories, USA) and nuclear counterstain (DAPI, 1:1000, D1306, Invitrogen, USA) diluted in blocking solution with shaking at 37°C for 3 days. The samples were then washed in 1X PBS three times with shaking at room temperature for 1 hour each. Next, samples were mounted in 1 mL syringes in agarose (2% (wt/wt), Agarose for ≧1kbp fragment, 01163-76, Nacalai USA, USA). Finally, the mounted samples were subjected to the RI-matching protocol described above.

### 1.5 Linear Expansion of Tissue

To assess tissue linear expansion as a result of clearing, the two-dimensional size of tissue samples was measured throughout the clearing protocol. Tissue pieces were placed on a glass dish with a ruler and imaged using a stereo microscope. Initial images of the tissue pieces were taken prior to starting the clearing protocol. Then, additional images were collected throughout the clearing protocol at the time points designated in **Table 3**. To quantify the images, tissue area was measured by thresholding images in FIJI (ImageJ)^21^. Linear expansion was calculated by normalizing the area of the cleared tissue with the area of the same tissue prior to starting the clearing protocol. Then the square root of this area size change was calculated.

**Table 3.** Linear Expansion Imaging Timeline.

| Day | Protocol Step | Imaging Notes |
| --- | --- | --- |
| 0 |  | Initial image |
| 1 | 1/2 CUBIC-L |  |
| 2 | CUBIC-L | Image before moving to CUBIC-L |
| 4 | Refresh CUBIC-L | Image |
| 6 | Refresh CUBIC-L | Image |
| 8 | Refresh CUBIC-L | Image |
| 10 | Refresh CUBIC-L | Image |
| 12 | Wash + Block | Image before and after washes |
| 13 | UEA and DAPI | Image before incubation |
| 16 | Wash + 1/2 CUBIC-R+ | Image before and after washes |
| 17 | CUBIC-R+ | Image before moving to CUBIC-R |
| 18 | Refresh CUBIC-R+ | Image |
| 20 | Image or store sample | Image |

### 1.6 Imaging Depth

To assess imaging depth before and after clearing, cleared and non-cleared tissue pieces were stained with PI. One z-stack of 500 μm with a z-step of 2.409 μm was acquired for each tissue piece on a Leica SP8 laser scanning confocal microscope. The laser power and gain were maintained across all tissue pieces. Imaging depth was calculated from contrast decay. Image contrast can be described by Equation 1 where 𝐼𝐼 corresponds to the grayscale value (or intensity) for each pixel, 𝐼_mean_ is the average intensity of the image, and 𝑛 is the total pixel count in the image. Accordingly, image contrast was acquired by measuring the standard deviation of the gray values for each image slice throughout the z-stack in FIJI (ImageJ) (Equation 2). Contrast values for each image were then normalized to the maximum contrast value in the image z-stack. Imaging depth was defined as the depth where contrast drops to ½ of the maximum contrast in the z-stack (i.e. the depth at which normalized contrast was equal to 0.5 given that the maximum normalized contrast was equal to 1).

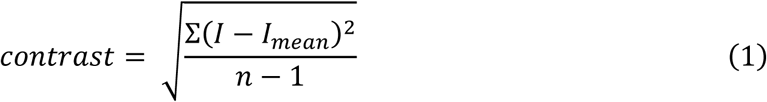

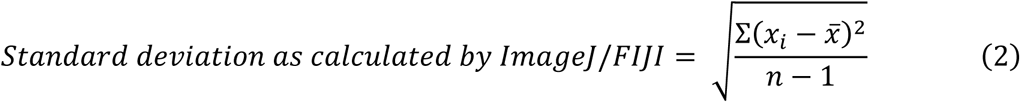

### 1.7 Evaluation of Follicle Density using Tissue Clearing

Follicle density was measured by staining cleared cortex tissue with ZP3 as described above. Tile scans (z-stack depth: 500 μm; z-step size: 2.5 μm) encompassing the full detectable tissue area were acquired using a Leica SP8 laser scanning confocal microscope. Follicles throughout the z-stacks were manually counted in the DAPI channel in FIJI (ImageJ), ensuring not to double-count follicles that spanned across multiple z-slices. Follicles were marked on a z-slice once the oocyte was fully visible. Follicle counts were then validated using the ZP3 channel. To calculate follicle density, tissue area was measured using FIJI (ImageJ). Specifically, starting from the top of the z-stack, tissue area was measured every 50 μm which corresponds to every 20^th^ z-slice. The volumes of these tissue sub-stacks were calculated by multiplying the tissue area with its respective height of 50 μm. Follicle density was then calculated for each tissue sub-stack as the total number of follicles in the sub-stack divided by the sub-stack tissue volume.

### 1.8 Evaluation of Follicle Density using Histology

Follicle density was assessed throughout a subset of fresh ovarian cortex samples which were fixed in Bouin’s fixative (Ricca Chemical, USA) at the time of tissue collection rather than cryopreserved. Samples were fixed at 4°C overnight and then washed in dH_2_O for 1 hour, 50% EtOH for 1 hour, 70% EtOH for 1 hour, and finally stored in 70% EtOH at 4°C. The fixed tissue samples were then embedded in paraffin at the University of Michigan Dental School Histology Core and serially sectioned at a thickness of 5 μm, producing four sections per slide for up to 50 slides per tissue sample. Every other slide was stained with hematoxylin and eosin (H&E) and imaged using a digital slide scanner (Leica Aperio AT2, Leica Biosystems, Germany) at 20× magnification at the University of Michigan Unit for Laboratory Animal Medicine Pathology Core.

Whole slide scans were imported into QuPath analysis software for manual follicle counting, as previously described^22^. Briefly, every 8^th^ tissue section was analyzed for follicles with 20 sections analyzed for donors A and B, 24 sections analyzed for donor C, and 25 sections analyzed for donor D. Analysis of every 8^th^ tissue section corresponded to a sampling spacing of 40 μm which prevented duplicate counts of a singular primordial follicle as they are approximately 40 µm in diameter^23^. As such, all visible primordial follicles were counted for each analyzed tissue section. Larger follicles were counted after comparing the preceding and subsequent tissue sections to prevent duplicate counts. To calculate tissue volume, tissue section area was multiplied by a tissue thickness of 20 μm rather than 40 μm to account for smaller primordial follicles that could have been omitted between every 8^th^ tissue section that was analyzed. Follicle density for each analyzed tissue section was then calculated as the total number of follicles in the tissue section divided by the tissue section volume.

### 1.9 Evaluation of Cortex Vasculature

Vasculature in human ovarian cortex samples was visualized by staining cleared tissue with UEA-1 as described above. Light sheet images (z-step size: 2.917 μm) were acquired using a Zeiss Lightsheet 7 at the University of Michigan BRCF Microscopy Core. Tile alignment was performed using Zeiss arivis Pro.

For quantitative analysis, image datasets were imported into Imaris. A region of interest (ROI) corresponding to ovarian cortex was defined based on tissue morphology, vascular features, and follicle stage composition. Within this ROI, vasculature was segmented by generating a surface with the Machine Learning Segmentation module. The resulting surface was used to create a mask of the UEA-1 signal which was subsequently analyzed with the Filament Tracer module to quantify branch points, total filament length, segment diameters, and vessel volumes. Total filament length and segment diameters were corrected for tissue linear expansion. Vessel length density and branch point density were calculated by using the expansion-adjusted ROI volume.

### 1.10 Statistical Analysis

All statistical analysis was performed in GraphPad Prism 10. Unless otherwise noted, data is expressed as mean ± standard deviation (mean ± SD). Error bars throughout figures are mean ± standard deviation per group.

## RESULTS

### 1.1. Adapting and Characterizing a Clearing Protocol for Human Ovarian Tissue

Among existing tissue clearing approaches, we selected CUBIC (clear, unobstructed brain imaging cocktails and computational analysis)^20,24^ for its versatility and proven performance across tissue types, including mouse ovaries^12^. To accommodate the larger size and denser stromal architecture of human ovarian tissue, the CUBIC protocol was adapted by extending the duration of its three major phases (**Figure 1A**): (1) delipidation and decolorization in CUBIC-L (∼10 days), (2) sample labeling (∼4–7 days depending on labeling method), and (3) refractive index (RI) matching in CUBIC-R (∼4 days). Implementation of this adapted protocol resulted in optically transparent ovarian cortex and medulla tissue (**Figure 1B**).

**Figure 1.**
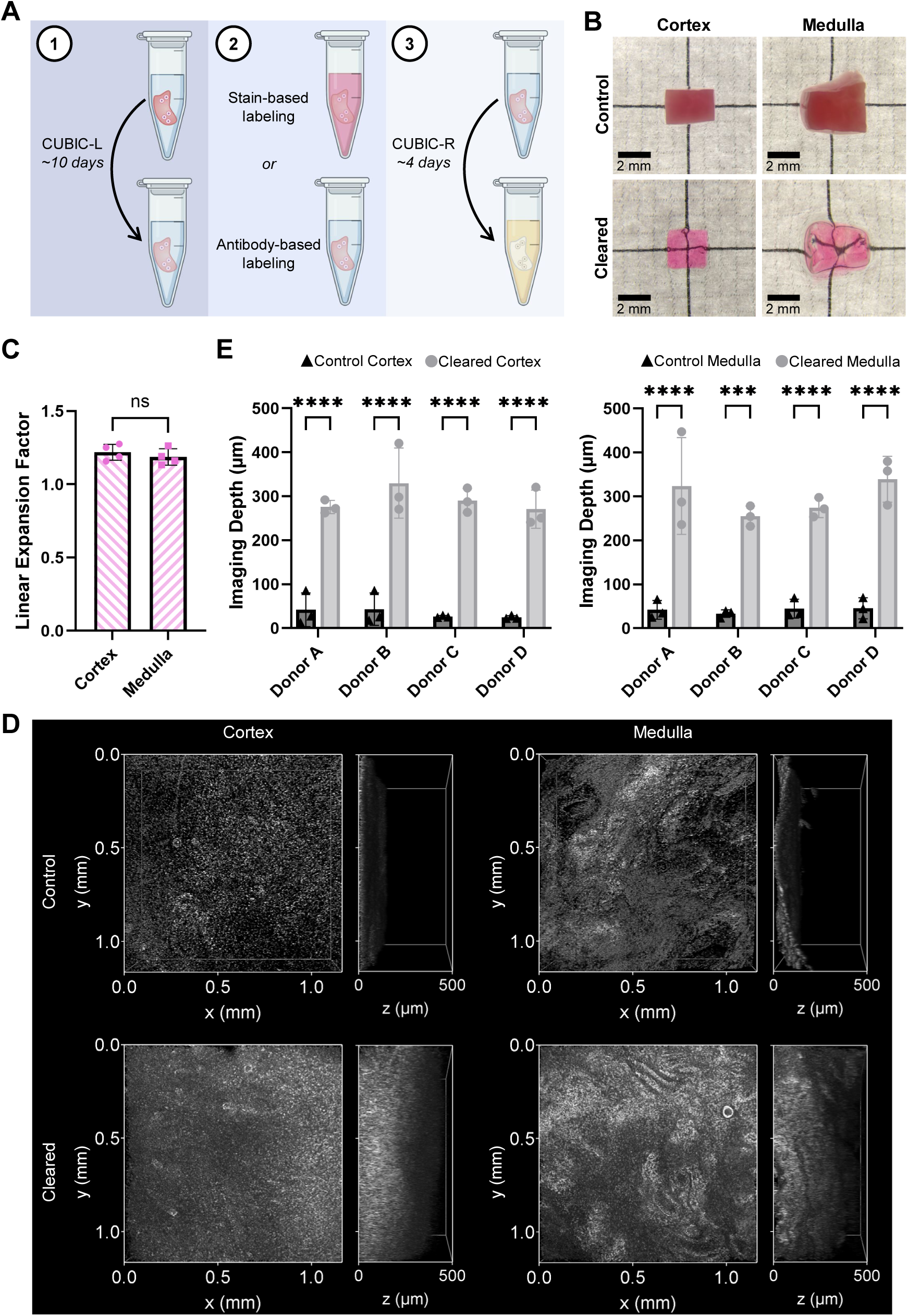
Adapted CUBIC Protocol Characterization for Human Ovarian Tissue. **(A)** Overview of the three major CUBIC protocol phases. **(B)** Cortex and medulla processed with the adapted CUBIC protocol become optically transparent, enabling visualization of the millimeter paper grid below. **(C)** Cleared cortex and medulla undergo similar expansion due to CUBIC clearing. Each data point represents the terminal linear expansion for a tissue piece (n = 4 tissue pieces per condition), error bars are mean ± standard deviation. Data analyzed by Mann-Whitney test, p < 0.05, ns = not significant. **(D)** Representative confocal images depicting the X-Y plane and Z imaging depth for control and cleared cortex and medulla. **(E)** CUBIC clearing increases imaging depth for both cortex and medulla across different tissue sources. Each data point represents the imaging depth for one tissue piece within the respective donor (n = 3 tissue pieces per donor), error bars are mean ± standard deviation. Data analyzed by two-way ANOVA with Šidák’s multiple comparisons test, p < 0.05, significant difference indicated as *p ≤ 0.05, **p ≤ 0.01, ***p ≤ 0.001, ****p ≤ 0.0001.

Because aqueous-based clearing protocols such as CUBIC can induce tissue expansion through hyper-hydration, changes in sample size over the course of clearing were quantified. By the end of the clearing protocol, both cortex and medulla exhibited a mean linear expansion factor of 1.2 relative to baseline starting size (**Figure 1C**), demonstrating consistent tissue swelling that was accounted for in downstream volumetric analyses.

To validate clearing performance, we compared imaging depth in control and cleared cortex and medulla samples from four donors using confocal microscopy (**Figure 1D**). Imaging depth was determined by contrast decay across z-stacks for each tissue piece (**Figure 1E**). Cleared cortex showed an approximately 6.5- to 11-fold increase in imaging depth relative to non-cleared controls and cleared medulla exhibited an approximately 6- to 8-fold increase, confirming that the adapted protocol substantially improved optical access in both tissue compartments.

### 1.2. Follicle Detection in Cleared Human Ovarian Cortex

We next investigated whether follicle density – a biologically and clinically relevant metric typically derived from two-dimensional serial histological sections – could be reliably quantified in cleared cortex. Cortical tissue was labeled for zona pellucida protein 3 (ZP3), which localizes to the extracellular matrix surrounding the oocyte. Cleared tissue was then imaged by confocal microscopy and follicles were identified morphologically in the DAPI channel with ZP3 labeling used for confirmation (**Figure 2A**).

**Figure 2.**
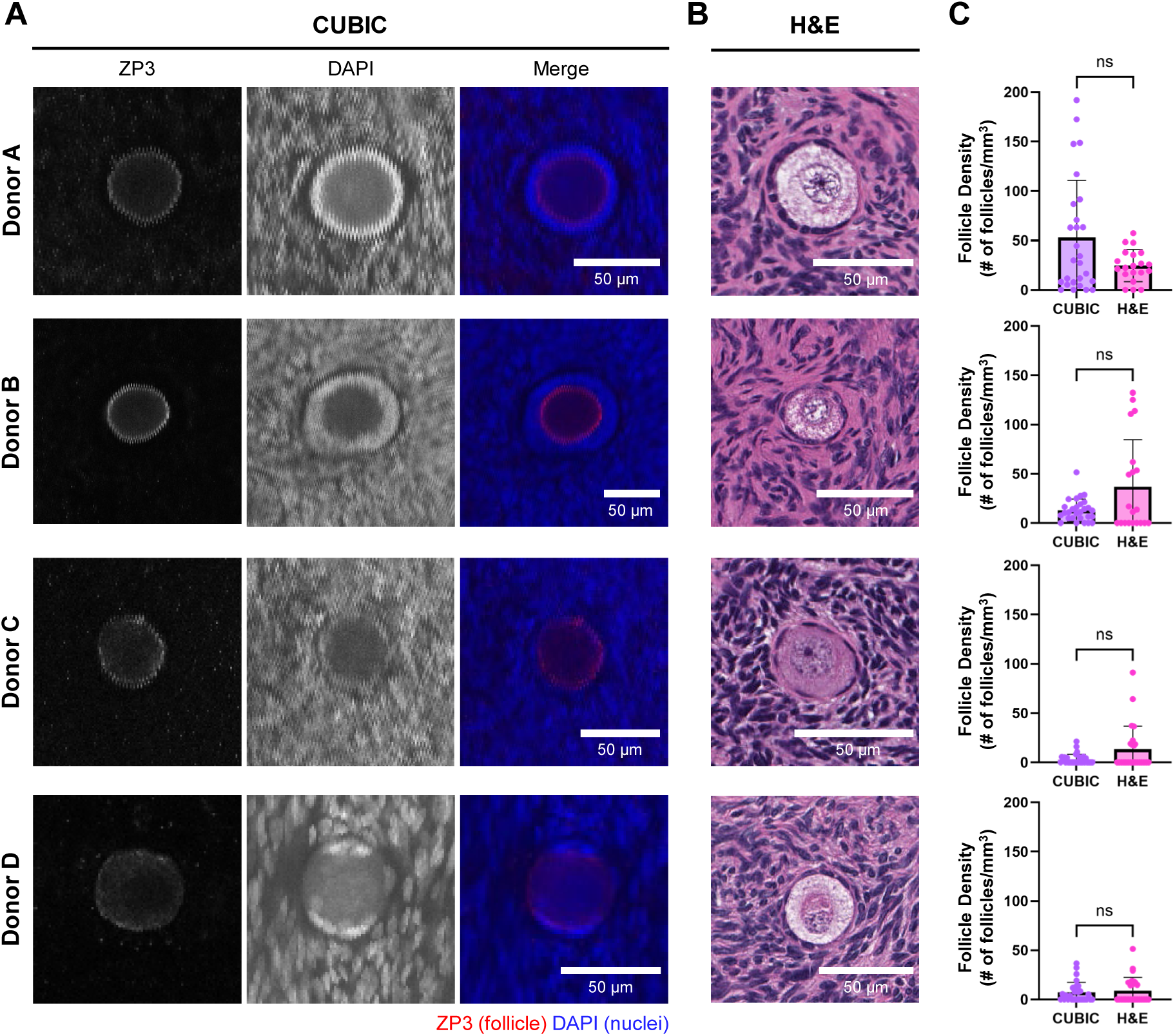
Follicle Detection and Quantification in Cleared Human Ovarian Cortex. **(A)** Representative confocal z-slices of follicles in cryopreserved cleared tissue depicting ZP3 and DAPI channels, and the merged images. **(B)** Representative histological H&E section images of follicles from fresh donor-matched tissue. **(C)** Comparison of follicle densities computed from CUBIC-cleared tissue and histological H&E images for each donor. Each data point represents the follicle density from a confocal z-stack substack (CUBIC) or a histological section (H&E), error bars are mean ± standard deviation. Data analyzed by Mann-Whitney test, p < 0.05, ns = not significant.

Follicle density was subsequently calculated within defined z-stack substacks for each tissue piece, enabling evaluation of follicle distribution throughout the imaged volume (**Figure S1, Table S1**). This analysis revealed heterogeneity both within and between donors. Donor A, for instance, showed notable variability across the analyzed tissue pieces with densities of 9 ± 9, 115 ± 58, and 35 ± 26 follicles/mm^3^. Across Donors B, C, and D, mean follicle densities ranged from 7–22, 1–7, and 1–16 follicles/mm^3^, respectively, further demonstrating the known intra-and inter-donor follicle distribution variability characteristic of human ovarian cortex.

Comparison of follicle densities from cleared tissue to follicle densities obtained using serial histological sections (**Figure 2B–C**) revealed no statistically significant difference across all donors, indicating that volumetric follicle quantification in cleared tissue was equivalent to standard histology while preserving the spatial context of follicles within the intact stromal environment.

### 1.3. Volumetric Characterization of Human Ovarian Cortical Vasculature

Given that follicle health and development are closely tied to stromal vascular support, particularly in the context of ovarian tissue cryopreservation and transplantation, we next examined native vasculature architecture within cleared human ovarian cortex. Tissue samples were labeled with UEA-1 lectin and imaged using light sheet microscopy, generating whole-tissue datasets that enabled visualization of vascular networks across intact tissue pieces (**Figure 3A**). For quantitative analysis, segmentation of vessel networks was restricted to sub-volumes consistent with cortical morphology (identified by the presence of small follicles, when possible) in order to minimize the inclusion of medullary regions. Within these regions of interest, vasculature was identified, traced (**Figure 3Ai**), and analyzed to extract morphometric parameters, including mean vessel diameter (**Figure 3Aii, Movie 1**), branch point density, length density, and volume fraction.

**Figure 3.**
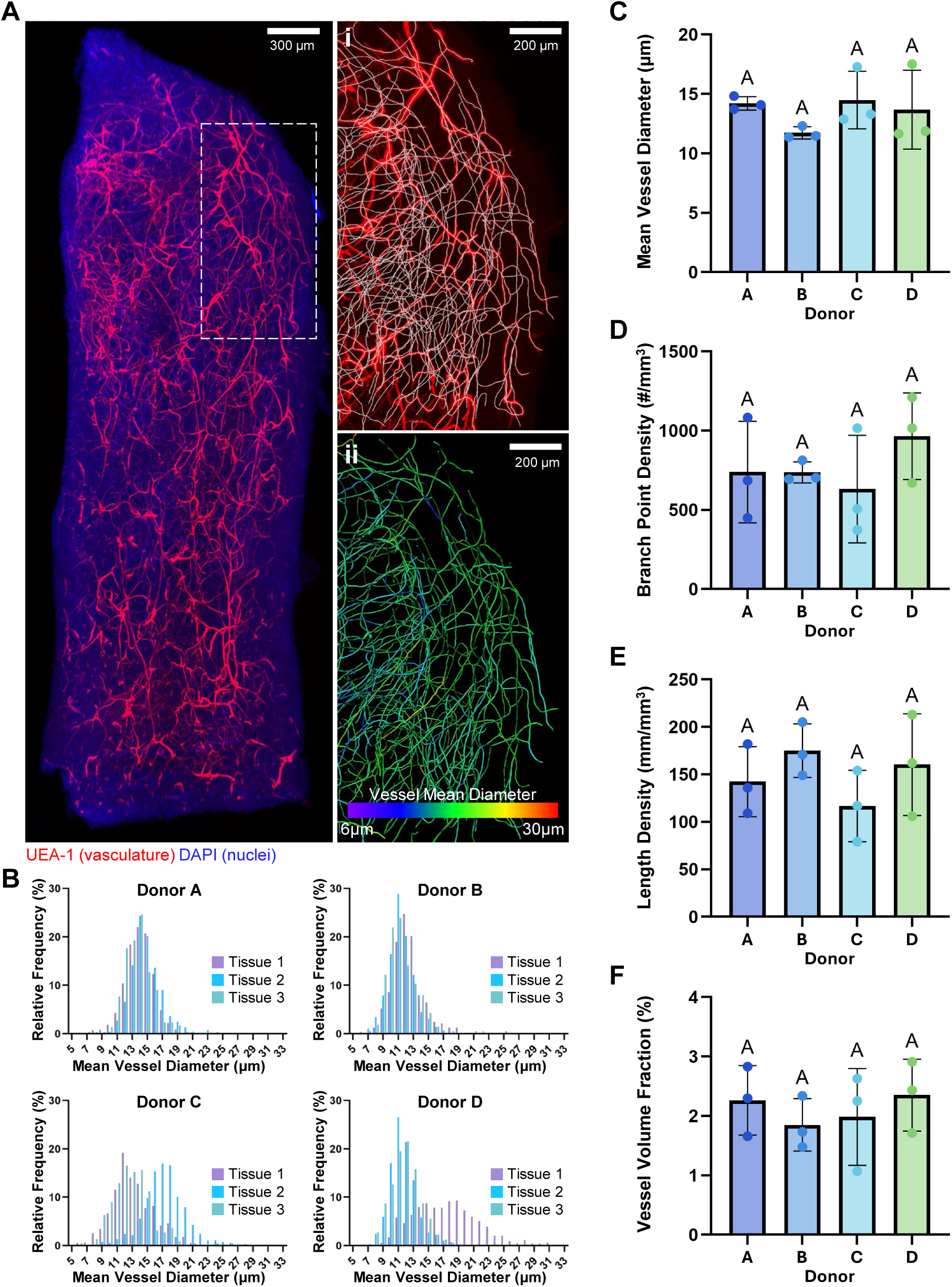
Vasculature Mapping in Cleared Human Ovarian Cortex. **(A)** Representative three-dimensional rendering of cortex vasculature from CUBIC-cleared and light sheet-imaged tissue with **(i)** showing vasculature tracing overlaid with the UEA-1 signal within an ROI and **(ii)** depicting the color-coded mean vessel diameters of the traced vasculature within the ROI. **(B)** Histograms of mean vessel diameter measurements for each donor. Within each histogram, data from three individual tissue pieces are shown and color-coded by tissue piece. For each tissue piece, blood vessels were automatically detected and segmented, and a mean diameter was calculated for every traced vessel. **(C)** Donor-level summary of mean vessel diameter. Each data point corresponds to one tissue piece and shows the mean of vessel mean diameters for the given tissue piece. Error bars are donor-level mean ± standard deviation. **(D)** Density of total vessel branch points detected within the analyzed ROI tissue volumes. **(E)** Density of total vessel length detected within the analyzed ROI tissue volumes. **(F)** Total vessel volume per tissue volume within analyzed ROI tissue volumes. Each data point represents the measurement for one tissue piece within the respective donor, error bars are mean ± standard deviation. Data analyzed by Kruskal-Wallis test with Dunn’s multiple comparisons test, p < 0.05, different letters indicate statistical significance.

Vessel diameter distributions ranged from 5 μm to 33 μm within individual tissue pieces (**Figure 3B**). When summarized for each donor, however, mean vessel diameters showed no statistically significant difference between donors (**Figure 3C**). Across donors, mean vessel diameter ranged from 12 µm to 14 µm, mean branch point density ranged from 632 points/mm^3^ to 965 points/mm^3^ (**Figure 3D**), mean vessel length density ranged from 117 mm/mm^3^ to 175 mm/mm^3^ (**Figure 3E**), and mean vessel volume fraction ranged from 1.9% to 2.3% (**Figure 3F**). Together, these findings demonstrate that tissue clearing combined with light sheet imaging enables quantitative, volumetric assessment of human ovarian cortical vasculature across intact tissue samples, capturing both local vascular heterogeneity and donor-level metrics.

### 1.4. Qualitative Spatial Relationships Between Follicles and Vasculature in Cortex

To illustrate spatial relationships between follicles and adjacent vasculature within the cleared human ovarian cortex, two-dimensional sections extracted from light sheet datasets were examined. In one region, a multilayer follicle was located next to a large vessel while a smaller nearby vessel appeared to partially wrap around the follicle (**Figure 4A**). Three-dimensional reconstruction of this region revealed the broader vascular network surrounding the follicle (**Movie 2**), demonstrating the extent and continuity of vessel segments rarely visualized in a single two-dimensional plane. Additional two-dimensional regions containing clusters of smaller follicles demonstrated variable degrees of vascular proximity (**Figure 4B**). In some regions, vessels were present within the surrounding stroma but remained separated from follicle boundaries, with the nearest vessel located approximately 100 μm (**Figure 4Bi**) or 50 μm (**Figure 4Bii**) from the closest follicle. In other regions, vascular structures approached more closely, located within 10 μm of the nearest follicle (**Figure 4Biii**), and even appeared to extend through the stroma between neighboring follicles (**Figure 4Biv**). These examples illustrate the range of follicle-vessel proximities present within a single tissue sample that are difficult to infer reliably from two-dimensional histological sections in the absence of volumetric imaging.

**Figure 4.**
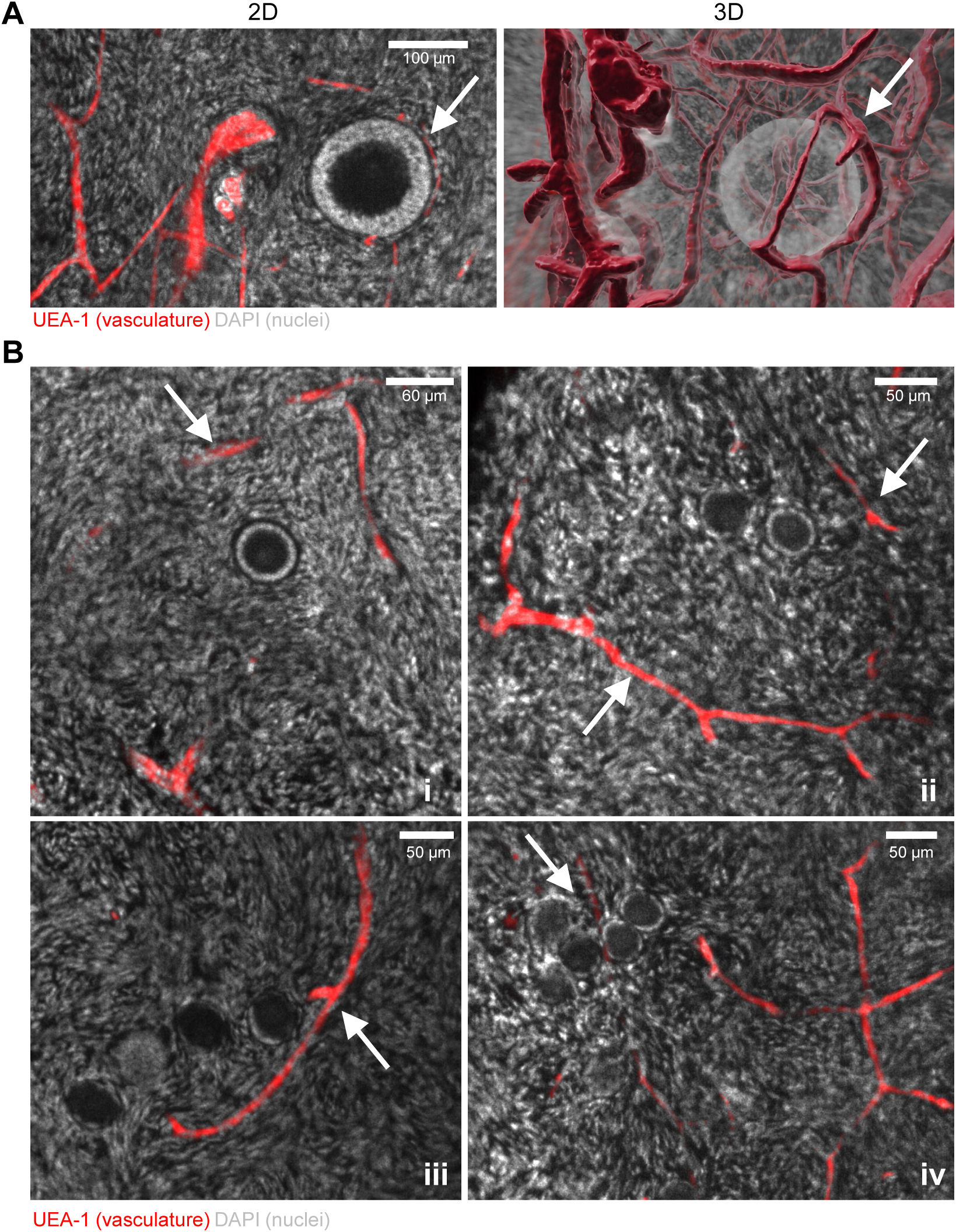
Location of Vasculature in Relation to Follicles. **(A)** Two-dimensional (2D) image of vasculature (white arrow) wrapping around a multilayer follicle. Vasculature traces from the previous light sheet vasculature analysis were overlaid with the follicle surface to better visualize the three-dimensional (3D) follicle-vessel spatial relationship. **(B)** Regions of cleared cortex with increasing proximity **(i–iv)** of vasculature (white arrows) to follicles.

### 1.5. Imaging Modality Considerations for Cleared Human Ovarian Tissue

To compare imaging approaches for cleared human ovarian tissue, the same cleared medulla sample was imaged using both confocal and light sheet microscopy. Light sheet microscopy achieved 8-fold greater imaging depth in one-sixth of the acquisition time relative to confocal microscopy (**Table 4**). These differences in imaging platforms translated into differences in structural visualization. In the confocal dataset, representative sections separated by 180 µm and 160 µm (**Figure 5A, Movie 3**) reconstructed only a limited portion of a large antral follicle. In contrast, light sheet sections separated by 485 µm and 367 µm (**Figure 5B, Movie 4**) resolved a substantially larger fraction of the follicle’s three-dimensional structure within the tissue. Despite these differences in volumetric coverage, both modalities still captured smaller adjacent follicles as well as surrounding granulosa and stromal cell populations within the medullary region (**Figure 5C**).

**Table 4.**
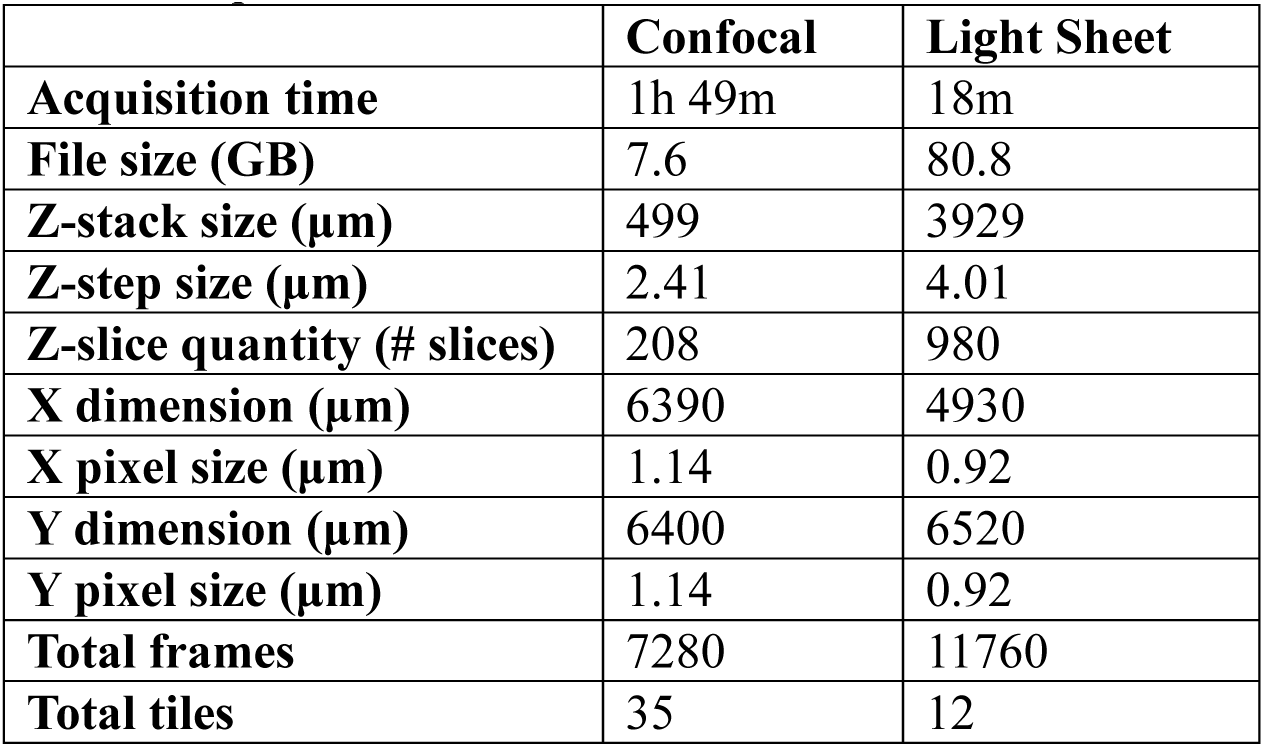
Acquisition Parameters and Dataset Characteristics for Cleared Medulla.

**Figure 5.**
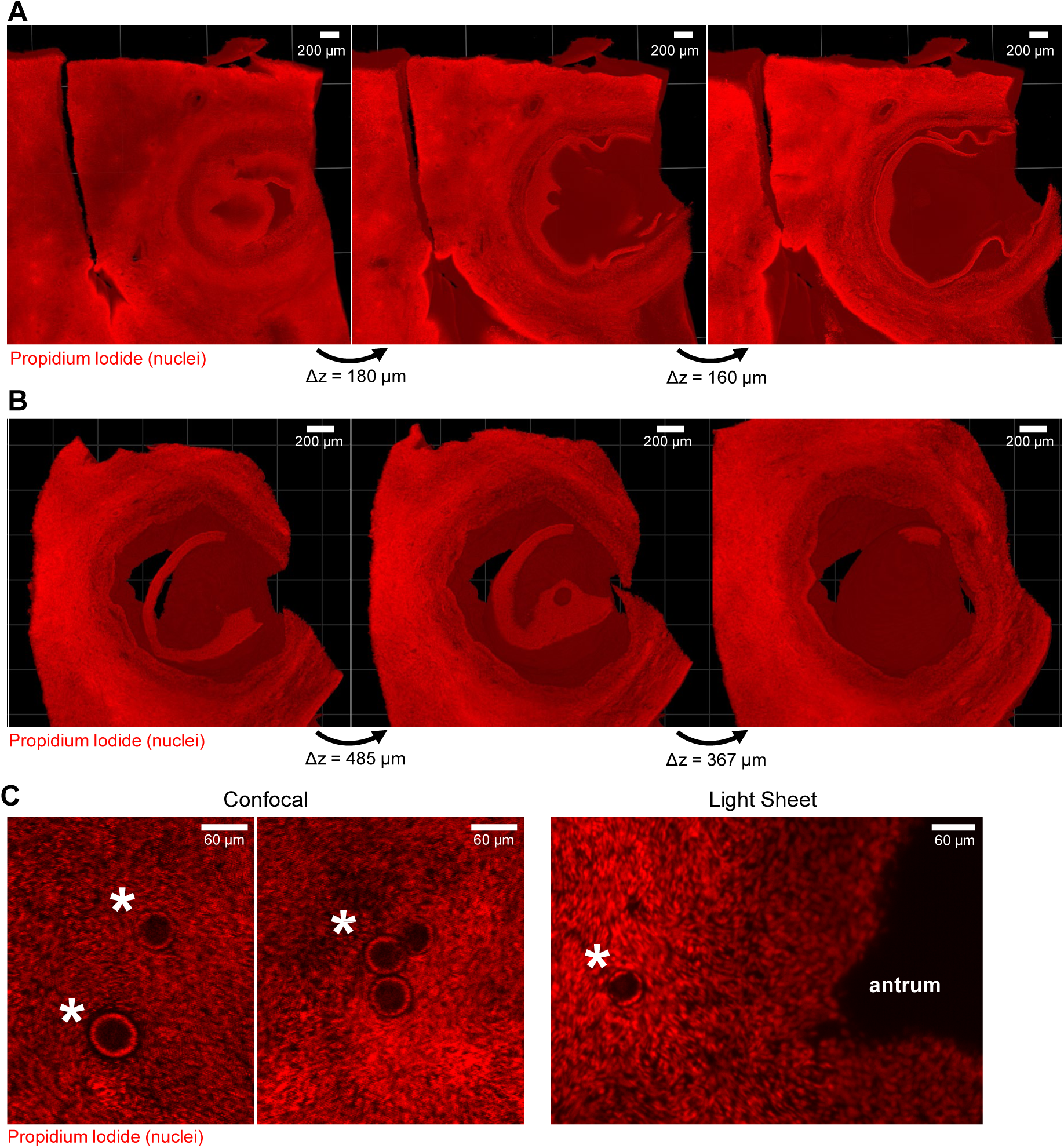
Comparison of Confocal and Light Sheet Imaging Modalities. **(A)** Sequential portions of a confocal z-stack depicting imaging depth capability in reference to an antral follicle structure. **(B)** Sequential portions of a light sheet z-stack of the same tissue piece depicting imaging depth capability in reference to the antral follicle structure. **(C)** Confocal z-slices depicting small follicles (asterisks) and surrounding stromal cells captured in the medulla sample. Light sheet z-slice depicting a small follicle (asterisk) and surrounding stromal cells adjacent to the follicle antrum in the medulla sample.

## DISCUSSION

This study establishes a reproducible tissue clearing and volumetric imaging workflow for cryopreserved human ovarian tissue that enables quantitative three-dimensional analysis of follicles and vasculature within intact tissue. By adapting a CUBIC-based protocol to accommodate the structural density and scale of human ovarian tissue, we demonstrate that ovarian volumetric readouts can be obtained without physical sample sectioning and that key metrics derived from cleared tissue are comparable to those generated using conventional histology. Together, these findings address a persistent methodological gap and expand the analytical toolkit available for studying human ovarian tissue in the context of fertility preservation and transplantation.

A central objective of this work was to determine whether follicle density measurements in cleared tissue were comparable to those obtained using the gold standard of histological sectioning. The agreement observed across all four donors supports the quantitative reliability of our workflow for follicle density assessment. This comparability, however, does not position tissue clearing as a replacement for histology. Histology remains the appropriate tool for routine follicle counting and staging, where detailed morphological assessment of individual follicles is the primary objective. In contrast, the value of tissue clearing lies in its capacity to preserve the spatial context of follicles within the surrounding stromal and vascular environment. By maintaining native tissue architecture, clearing enables simultaneous visualization and analysis of multiple tissue compartments in three dimensions, providing complementary information that cannot be obtained through serial histological sectioning alone.

Beyond follicle readouts, light sheet imaging of lectin-labeled cleared cortex enabled quantitative characterization of vascular architecture across intact tissues. The absence of significant donor-to-donor differences in vessel diameter, branch point density, length density, and volume fraction, despite within-tissue heterogeneity, suggests that baseline cortical vascular organization is relatively conserved across similarly aged individuals, a finding with potential implications for interpreting vascular remodeling after ovarian tissue transplantation. An additional strength of this workflow is the ability to visualize follicles and vasculature together within the same tissue volume. Follicle-vessel proximity varied markedly across tissues, ranging from distances on the order of 100 μm to less than 10 μm. While it remains to be determined whether these patterns reflect active vascular recruitment or passive architectural heterogeneity, volumetric imaging uniquely enables these spatial relationships to be observed in their native three-dimensional context.

Comparison of confocal and light sheet microscopy on the same cleared tissue highlighted important practical considerations for implementation of our workflow. Light sheet microscopy provided substantially greater volumetric coverage in a shorter acquisition time, facilitating the reconstruction of large follicular structures. The resulting datasets, however, were an order of magnitude larger and required dedicated computational resources. Confocal microscopy, while more limited in imaging depth, remains broadly accessible and produces datasets that are easier to manage using standard software. For applications focused on follicle density within cortical tissue, confocal imaging is adequate. In contrast, light sheet microscopy is preferable for studies requiring whole-tissue vascular network analysis or reconstruction of large follicular structures. As such, selection of imaging modality should be guided by the biological question and available infrastructure.

Several limitations should be considered. The donor cohort spanned a narrow age range, precluding assessment of age-associated changes in follicle density or vascular organization – a question of considerable relevance given the age-dependence of both the follicle reserve and ovarian vascularity^11,25^. Quantitative analysis was also restricted to the ovarian cortex, which is the most clinically relevant tissue for fertility preservation, but extending our workflow to the ovarian medulla would broaden its utility for studying follicle maturation and vascular remodeling across the full organ. Additionally, follicle and vascular metrics were quantified from separately labeled tissue pieces which prevented direct, systematic measurements of follicle-vessel proximity. Although follicles and vessels could be visually identified within the light sheet datasets, simultaneous labeling of these structures would enable robust and automated quantification of these spatial relationships. Incorporating dual-labeling and co-registration approaches will therefore be an important next step in this workflow. Finally, stromal compartments, including fibroblasts, smooth muscle cells, and extracellular matrix components, were not characterized in this study but inclusion of such stromal markers would further expand the mechanistic utility of our workflow.

In summary, this work provides a validated, reproducible framework for volumetric analysis of follicles and cortical vasculature in intact cryopreserved human ovarian tissue. Our approach enables quantitative analysis of these features while preserving the three-dimensional relationship relationships between follicles, vasculature, and the surrounding stromal microenvironment, thereby complementing traditional histology with spatial information that cannot be captured solely through serial sectioning. As fertility preservation strategies evolve towards understanding the interaction between follicle survival and vascular remodeling within the ovarian stroma, tools that enable quantitative, three-dimensional analysis of these relationships will become essential. The platform described here provides a foundation for these future investigations.

## SUPPLEMENTARY MATERIAL

**Figure S1.**
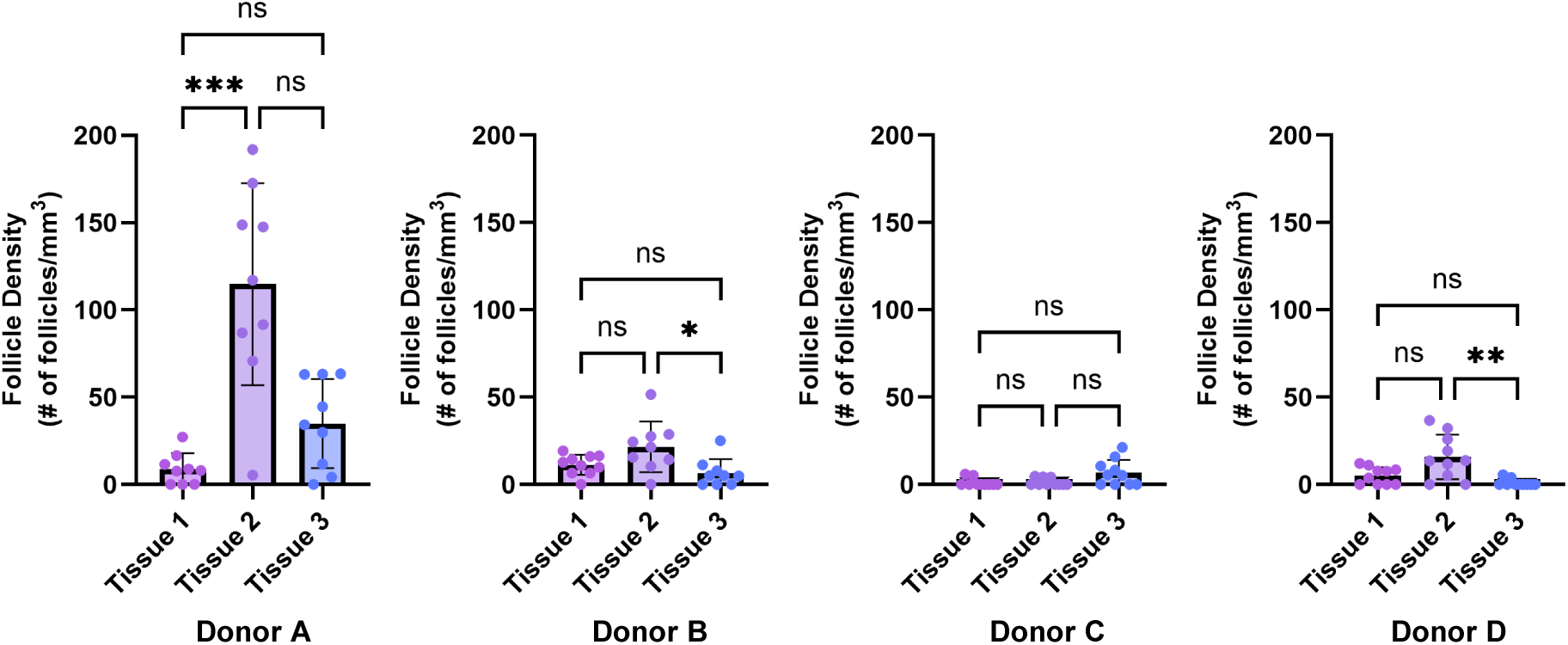
Intra-Donor Follicle Density Heterogeneity in Cleared Tissue. Follicle densities across each of the three analyzed tissue pieces for Donors A–D. Each data point represents the follicle density within one z-stack substack (9-10 substacks per tissue piece), error bars are mean ± standard deviation per tissue piece. Data analyzed by Kruskal-Wallis test with Dunn’s multiple comparisons test, p < 0.05, significant differences indicated as *p ≤ 0.05, **p ≤ 0.01, ***p ≤ 0.001.

**Table S1.**
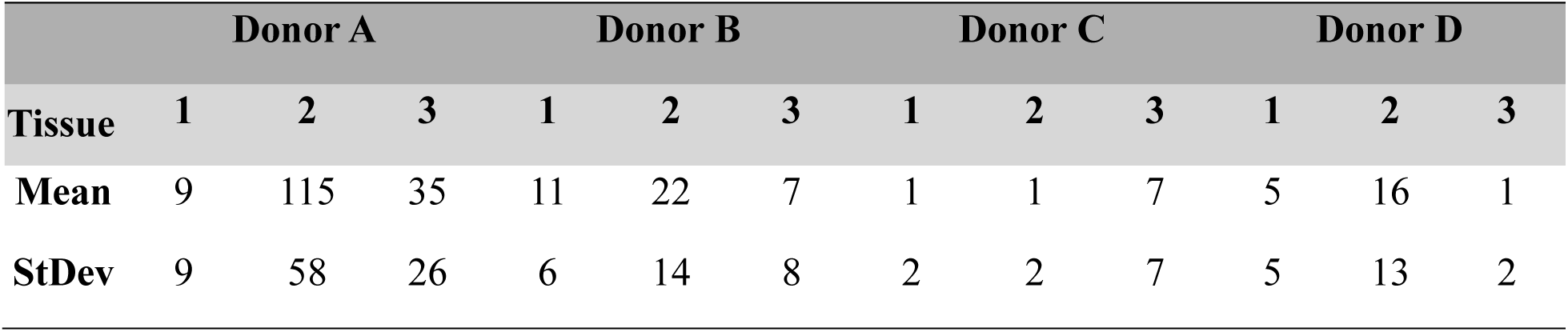
Tissue-Specific Follicle Densities in Cleared Tissue. Mean and standard deviation for follicle densities across each of the three analyzed tissue pieces for Donors A–D. Follicle density is expressed as number of follicles per mm^3^.

|  | Donor A |  |  | Donor B |  |  | Donor C |  |  | Donor D |  |  |
| --- | --- | --- | --- | --- | --- | --- | --- | --- | --- | --- | --- | --- |
| Tissue | 1 | 2 | 3 | 1 | 2 | 3 | 1 | 2 | 3 | 1 | 2 | 3 |
| Mean | 9 | 115 | 35 | 11 | 22 | 7 | 1 | 1 | 7 | 5 | 16 | 1 |
| StDev | 9 | 58 | 26 | 6 | 14 | 8 | 2 | 2 | 7 | 5 | 13 | 2 |

## MOVIE STILLS

**Movie 1.**
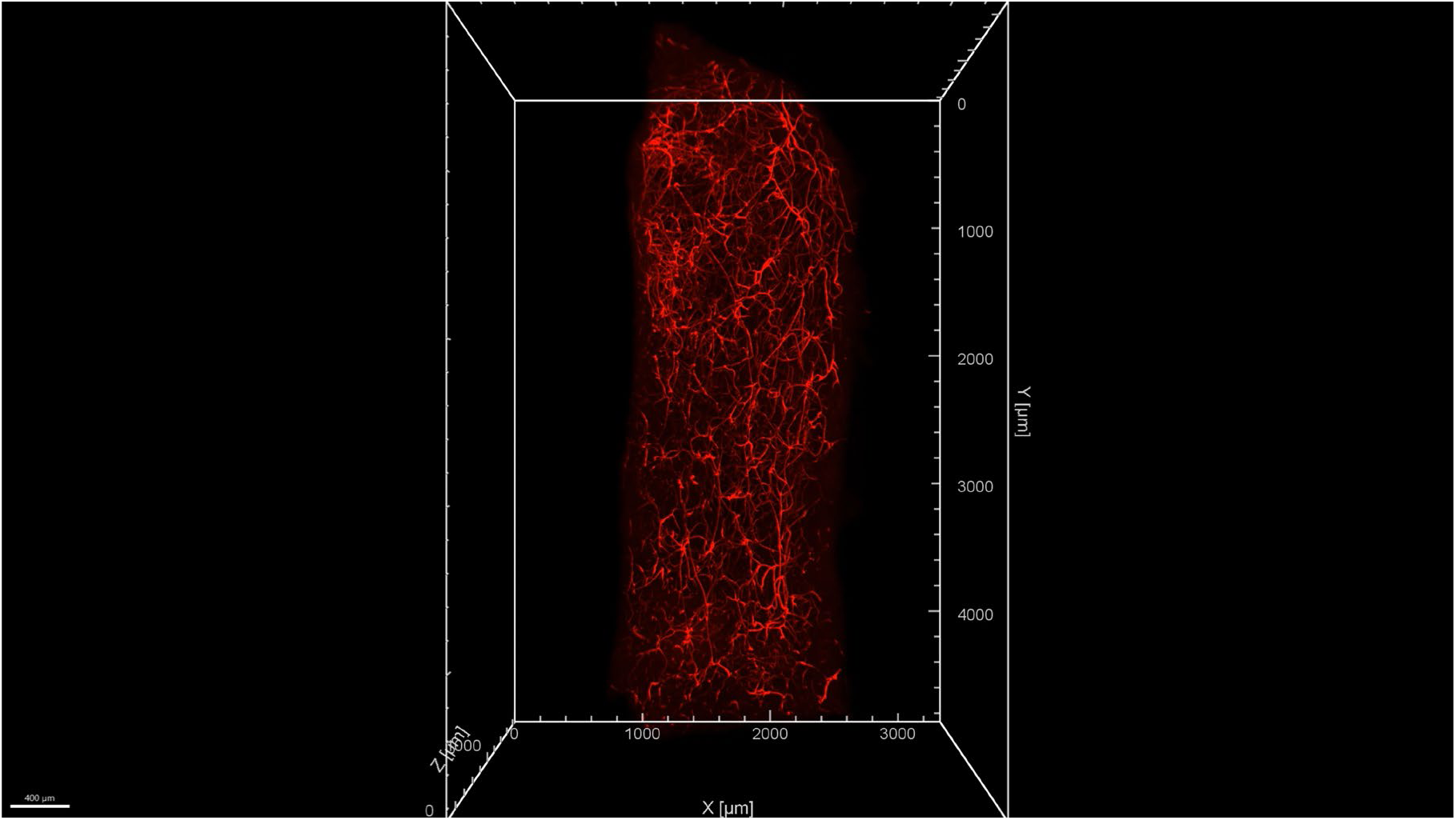
Three-dimensional visualization of vascular networks (red) across an intact human ovarian cortex sample combined with vasculature tracing (grey) and mean vessel diameter analysis (heatmap) performed in Imaris.

**Movie 2.**
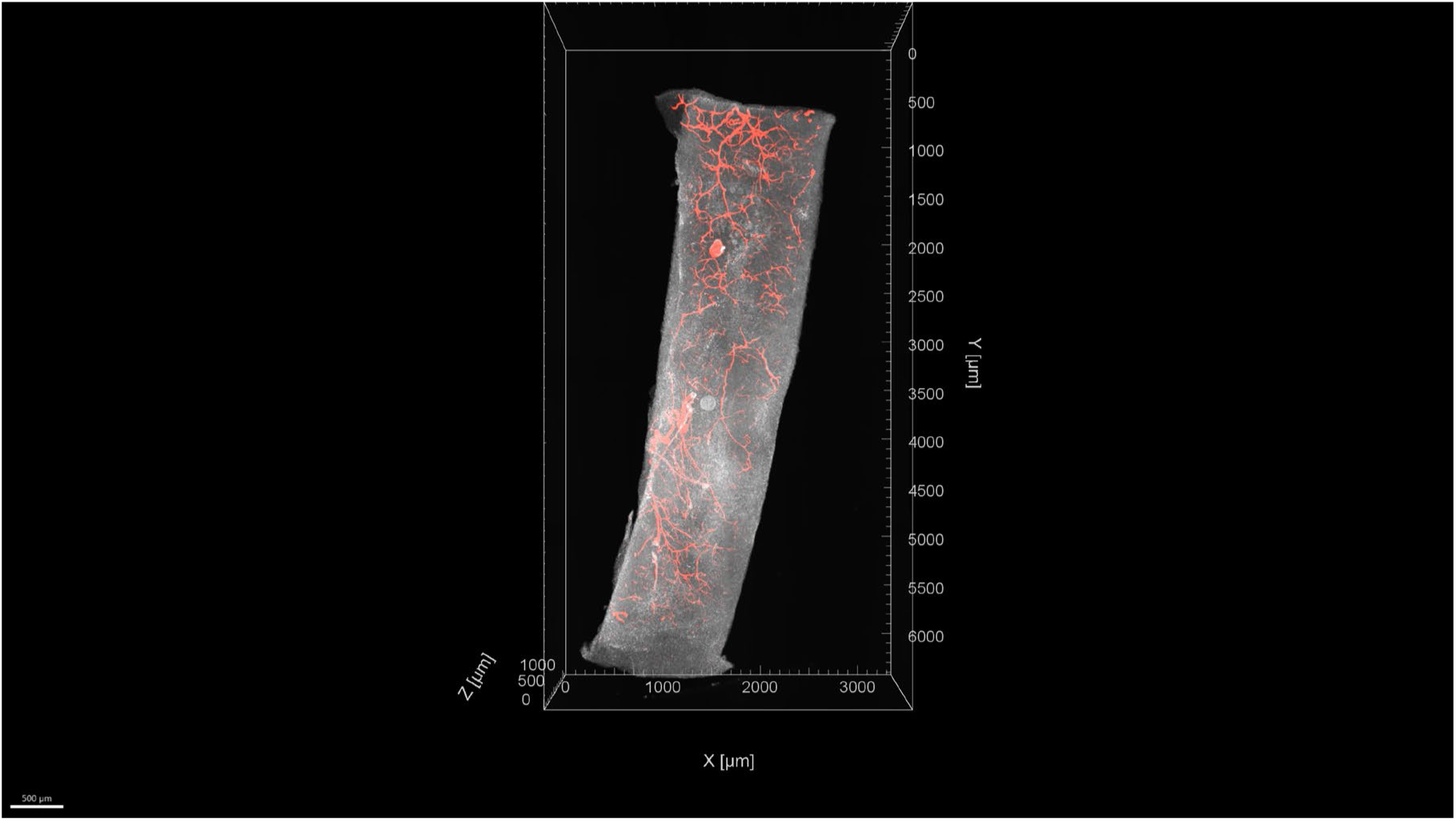
Light sheet imaging captures the three-dimensional relationship between vasculature (red) and follicle (grey).

**Movie 3.**
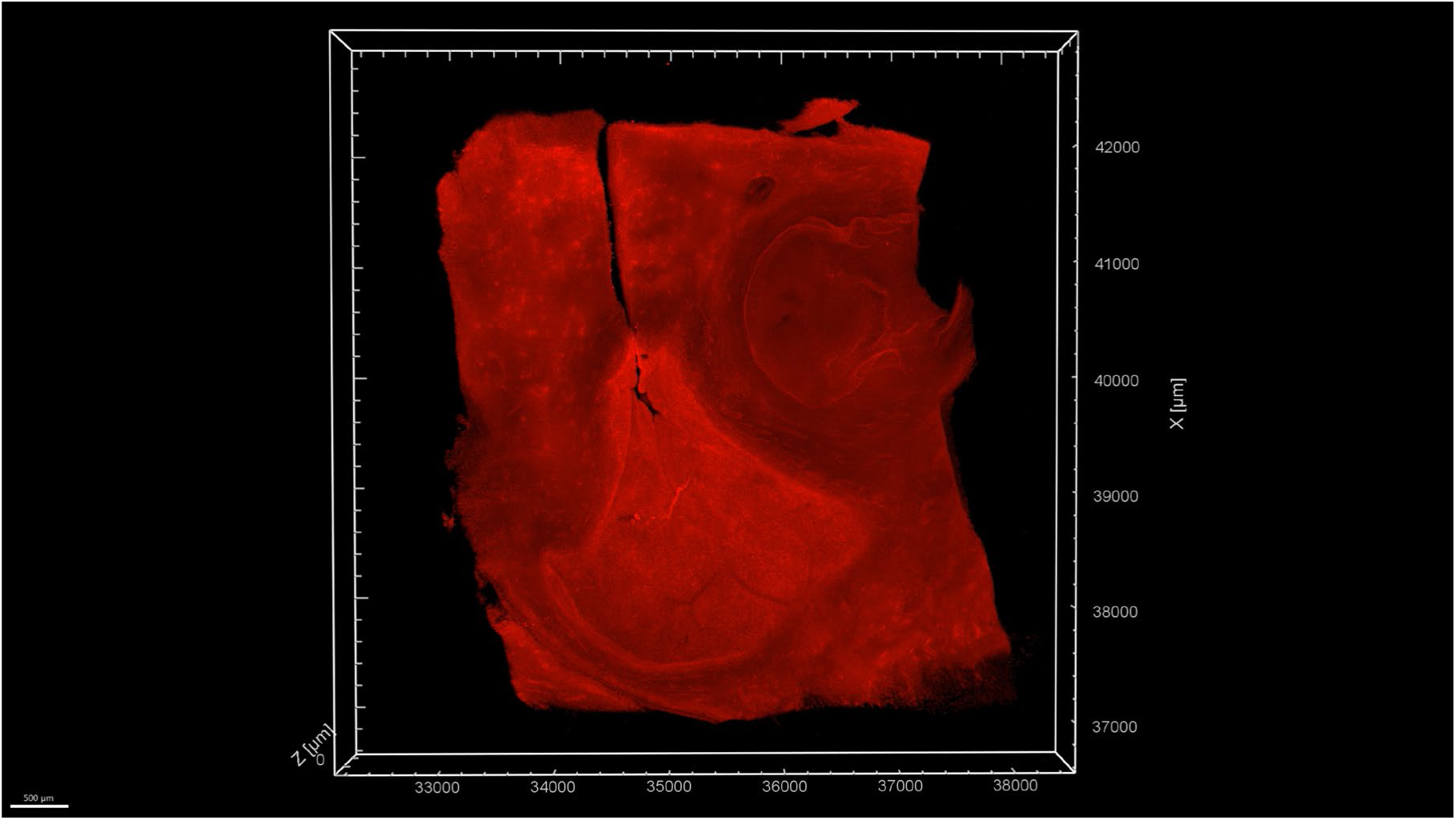
Confocal microscopy captures a portion of a large antral follicle.

**Movie 4.**
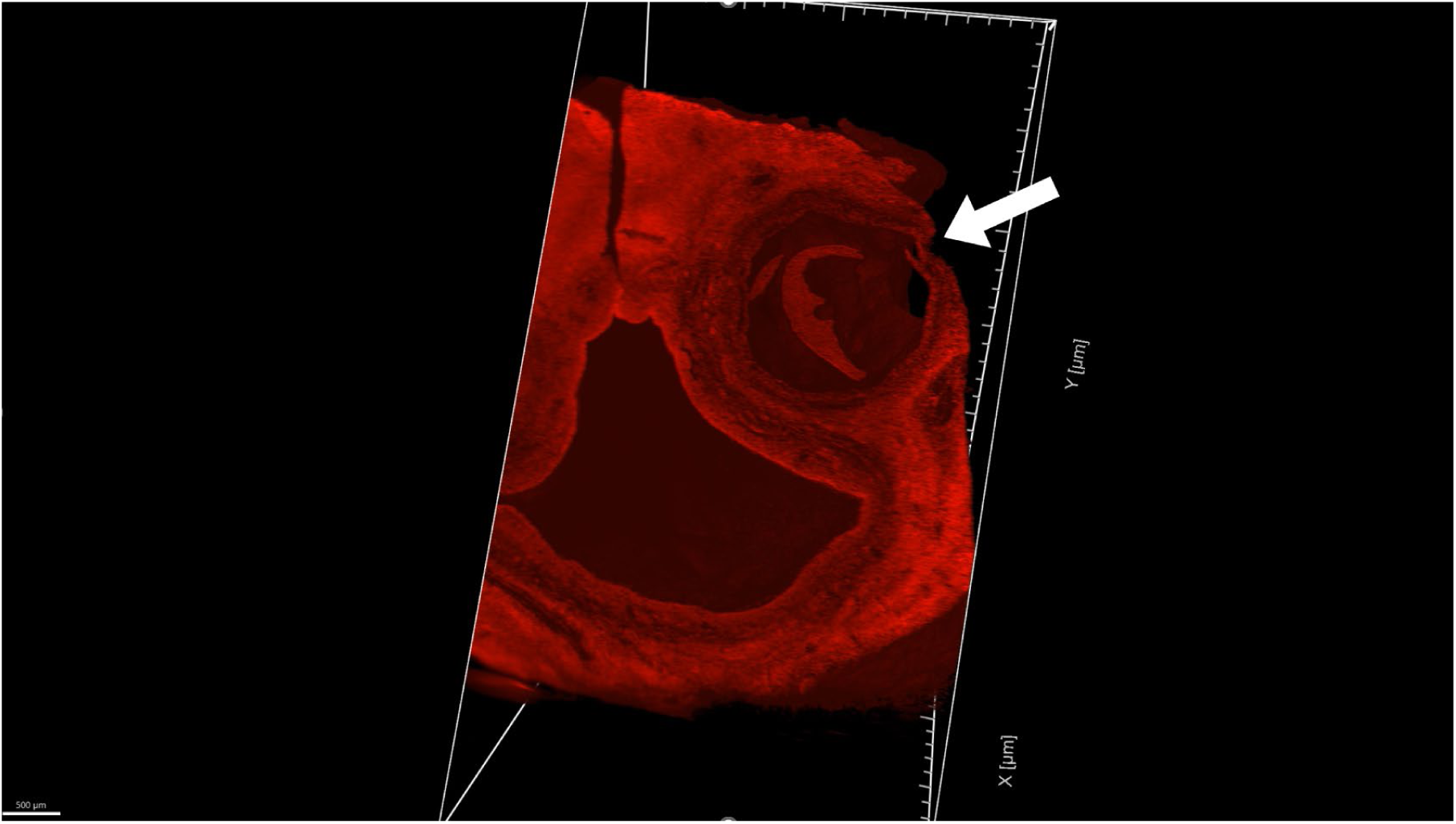
Light sheet microscopy resolves the entire large antral follicle structure (white arrow).

## ACKNOWLEDGEMENTS

We thank the members of the Shikanov and Baker laboratories for scientific discussions. We would like to acknowledge the University of Michigan Dental School Histology Core and the Unit for Laboratory Animal Medicine Pathology Core for histological processing and whole slide scanning, respectively. We would also like to acknowledge the University of Michigan BRCF Microscopy Core staff Binyamin Jacobovitz for training and advice on light sheet microscopy. Illustrative schematics were created with BioRender. Lastly, we thank all human organ donors provided by our partnership with the International Institute for the Advancement of Medicine (IIAM).

## AUTHOR CONTRIBUTIONS

Conceptualization and Ideation: DIP, BMB, AS

Study Design: DIP, BMB, AS

Sample Clearing and Processing: DIP, CEF

Cleared Sample Image Acquisition: DIP

Cleared Sample Image Analysis: DIP, CEF, MAJ

Histological Sample Analysis: JHM, VJ

Manuscript Drafting: DIP, AS

Review, and Editing: DIP, BMB, AS

## GENERATIVE AI TOOLS

During the preparation of this work, the authors used the University of Michigan GPT-4.1 in order to proofread sentences and paragraphs for spelling, grammar, and clarity. The platform was used for language refinement and not for content generation. After using this tool, the authors reviewed and edited the content and take full responsibility for the content of the published article.

